# A mechanistic link between cellular trade-offs, gene expression and growth

**DOI:** 10.1101/014787

**Authors:** Andrea Y. Weiβe, Diego A. Oyarzún, Vincent Danos, Peter S. Swain

## Abstract

Intracellular processes rarely work in isolation but continually, interact with the rest of the cell. In microbes, for example, we now know that gene expression across the whole genome typically changes with growth rate. The mechanisms driving such global regulation, however, are not well understood. Here we consider three trade-offs that because of limitations in levels of cellular energy, free ribosomes, and proteins are faced by all living cells and construct a mechanistic model that comprises these trade-offs. Our model couples gene expression with growth rate and growth rate with a growing population of cells. We show that the model recovers Monod's law for the growth of microbes and two other empirical relationships connecting growth rate to the mass fraction of ribosomes. Further, we can explain growth related effects in dosage compensation by paralogs and predict host-circuit interactions in synthetic biology. Simulating competitions between strains, we find that the regulation of metabolic pathways may have evolved not to match expression of enzymes to levels of extracellular substrates in changing environments but rather to balance a trade-off between exploiting one type of nutrient over another. Although coarse-grained, the trade-offs that the model embodies are fundamental, and, as such, our modelling framework has potentially wide
 application, including in both biotechnology and medicine.

---

Intracellular processes rarely work in isolation but continually interact with the rest of the cell. Yet often we study cellular processes with the implicit assumption that the remainder of the cell can either be ignored or provides a constant, background environment. Work in both systems and synthetic biology is, however, showing that this assumption is weak, at best. In microbes, growth rate can affect the expression both of single genes [1, 2] and across the entire genome [3, 4, 5, 6]. Specific control by transcription factors appears to be complemented by global, unspecific regulation that reflects the physiological state of the cell [5, 6, 7]. Correspondingly, progress in synthetic biology is limited by two-way interactions between synthetic circuits and the host cell that cannot be designed away [8, 9].

These phenomena are thought to arise from trade-offs where commitment of a finite intracellular resource to one response necessarily reduces the commitment of that resource to another response. A trade-off in the allocation of ribosomes has been suggested to underlie global gene regulation [2,5]. Similarly, depletion of finite resources and competition for cellular processes is thought to explain the failure of some synthetic circuits [8]. Such circuits ′load′ the host cell, which can induce physiological responses that further degrade the function of the circuit [10]. Our understanding of such trade-offs, however, is mostly phenomenological.

Here we take an alternative approach and ask what new insight can be gained from a minimal mechanistic model that captures these trade-offs. We focus on three trade-offs that can be considered universal in the sense that they are experienced by all living cells: (i) finite levels of cellular energy so that launching a new biochemical process reduces the activities of others; (ii) finite levels of ribosomes so that translating a new type of mRNA reduces translation of all other mRNAs; and (iii) a finite proteome, or cell mass, so that expressing a new type of protein reduces levels of other types. Reduced demand on any of these finite resources will, correspondingly, free that resource for other intracellular processes.

We develop a mechanistic cellular model built around these three trade-offs. The model predicts allocation of the proteome, energy turnover, and physiological phenotypes, such as growth rate, from specifications made at the level of genotype, and thus connects molecular mechanisms to cellular behaviour. A whole-cell model has been proposed as one way to make such predictions [11], but its level of detail may sometimes obscure the core biochemistry that underlies the observed phenotypes and potentially complicates further analyses. We instead adopt a complementary coarse-grained approach [12,13, 14] and try to find minimal descriptions that highlight the mechanisms generating the *in silico* phenotypes we observe. In contrast to other approaches [13, 14], we emphasize that we do not optimize either growth rate or any other physiological variable.

With only these trade-offs, we can derive fundamental properties of microbial growth [15, 16] and potentially explain diverse phenomena such as gene dosage compensation [17] and host effects on the performance of synthetic circuits. Our mechanistic framework can be extended to include, for example, signal transduction and population-scale effects. Using such an extension, we study the evolutionary benefits of gene regulation and find that transcriptional regulation of metabolic pathways may have evolved to balance the uptake of different nutrients rather than to tune levels of enzymes to match the extracellular availability of their substrates in changing environments.

## Results

### Using trade-offs to construct a mechanistic single-cell model

Our model implements trade-offs faced by cells by considering two core biochemical processes: gene expression and nutrient import and metabolism (Fig. 1A). To focus on the effects of the trade-offs, the model is a deterministic system of ordinary differential equations, each one describing the rate of change of the numbers of molecules per cell of a particular intracellular chemical species. Throughout, we work with numbers of molecules rather than concentrations and, for simplicity, do not explicitly model changes in cell volume. Details of the model are given in the SI (SI Appendix, §S.1).

**Finite energy:** The first trade-off that we include is the finite size of the pool of intracellular levels of energy. We consider a generic form of energy, denoted a, that includes all intracellular molecules used to fuel molecular synthesis, such as ATP and NADPH. The environment contains a single nutrient, *s*, that once internalized (and then denoted *s_i_*) can be metabolized. One molecule of *s* yields *n_s_* molecules of *a*. If *e_t_* denotes the enzyme that transports *s* into the cell and *e_m_* denotes the enzyme that metabolizes *s_i_* into *a*, then the dynamics of *s_i_* obey

**Figure 1.**
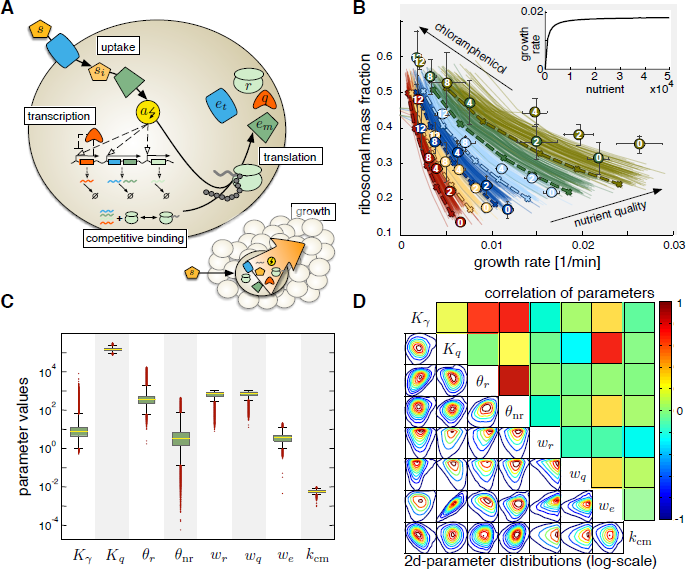
– A mechanistic cell model that recovers the laws of microbial growth. A. Schematic of the model. Enzymes (shown in blue and dark green) import and metabolize an extracellular nutrient (shown in orange), which yields energy (yellow). Transcription of all genes depends on energy (dashed arrows). mRNA molecules compete for ribosomes (light green). The overall rate of translation determines the rate of growth (lower right). We model three classes of proteins: ribosomes, enzymes and other house-keeping proteins, *q* (red). *B*. The model fits the data from Scott *et al.* [2] that empirically demonstrate two of the growth relations. Growth rate is changed by either changing the quality of nutrients (dots of the same color indicate the same extracellular media) or by adding chloramphenicol, a drug that inhibits translation (numbers within dots indicate the concentration in μM). Solid lines show the fits from 100 parameter sets randomly drawn from the posterior distribution; dashed lines are the fit given by the modes of the marginal posterior distributions, which we used in subsequent simulations. Inset: Varying the amount of external nutrient, the model reproduces Monod's growth law. *C*. The posterior probability distributions of the parameters show no fine-tuning. Box plots indicate the median, the 25%, and 75% quantiles with outliers in red. The distributions span several orders of magnitudes (except those of *K_q_ and k_cm_*) indicating that the parameter fit is robust. *D*. Statistical dependencies between parameter values show that a few pairs of parameters are strongly correlated. Lower triangle: Pairwise posterior distributions. Upper triangle: correlation coefficient.

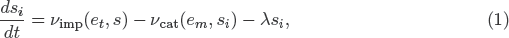

where the rates of import, *v_imp_*, and of metabolism, *v_cat_*, both have a Michaelis-Menten form. The growth rate is denoted by λ, and all intracellular species are diluted at a rate λ because of partitioning of molecules to daughter cells at division.

For both *E. coli* and *S. cerevisiae*, the two best studied microbes, translation dominates the consumption of cellular energy [18,19,20], and, in the spirit of a minimal model, we therefore neglect other energy-consuming processes. If each translational elongation step consumes one unit of a, then the amount consumed during the translation of a protein *x* is proportional to its length *n_x_*. Letting *v_x_* denote the translation rate for protein *x*, we can describe the overall turnover of energy by

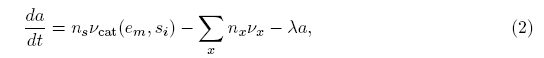

where the sum over x is over all types of protein in the cell. We see that energy is created by metabolizing *s_i_* and lost through translation and dilution. The effective rate of translational elongation obeys

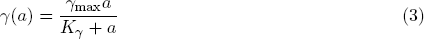

if an equal amount of energy is consumed for the addition of each amino acid to the growing peptide chain (see SI Appendix, §S.1.2.3). Here γ_max_ is the maximal elongation rate and *K_γ_* is the threshold amount of energy where elongation is half-maximal. Using *c_x_* to denote the complex between a ribosome and the mRNA for protein *x*, then the translation rate for *x* is

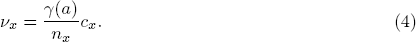

It is through the sum in Eq. 2 and the energy dependence of Eq. 4 that the first trade-off is implemented. Translation of each mRNA consumes *a*, and levels of *a* determine the rate of translation of all mRNAs.

**Finite ribosomes:** The second trade-off results from the finite pool of intracellular ribosomes. To include this trade-off, we explicitly model the competition between mRNAs for binding free ribosomes. Let *r* denote the number of free ribosomes. Let *k_b_* and *k_u_* denote the rates of binding and unbinding of a ribosome to mRNA (assumed identical for all mRNAs) and let the mRNA for a protein *x* be *m_x_*, then

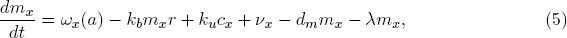

with *ω_x_ (a)* being the rate of transcription. The rate *d_m_* is the rate of degradation of all mRNAs (assumed equal for simplicity). Similarly, for the ribosome-mRNA complex, we have

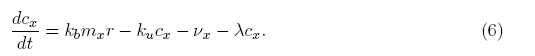

Translation, by releasing *m_x_* from *c_x_*, contributes a positive term to Eq. 5 and a negative term to Eq. 6. Again, in the spirit of a minimal model, we do not include polysomes but assume an mRNA can bind only one ribosome. The equation for free ribosomes is

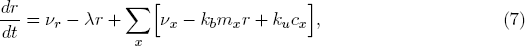

where the sum over all proteins, including ribosomes, again implements the trade-off.

Although we neglect the contribution of processes other than translation to the consumption of energy, we do model transcription as dependent on levels of energy because transcription must cease when all energy is lost. Analogous to our model of translation, Eq. 3, if each transcriptional elongation step uses a fixed (though assumed negligible) amount of energy, it follows that the transcription rate for a gene *x* takes the form:

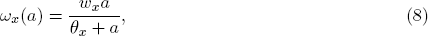

where *ω_x_* is the maximal transcription rate and *θ_x_* is the threshold amount of energy at which transcription is half-maximal. We note that *ω_x_* is determined by the copy number, induction level, and length of gene *x*. Eq. 8 holds too for ribosomes. Although ribosomes are ribonucleoproteins, we ignore such complexity and consider only the expression of their protein component because only the protein component is necessary to implement the tradeoffs.

Besides ribosomes, we include other house-keeping proteins, such as cytoskeletal proteins. Denoting these proteins by *q*, we assume their transcription to be negatively auto-regulated to maintain stable levels across different growth conditions [1,2]: 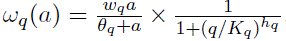.

**Finite proteome:** Finally, we include the third trade-off, the finiteness of the proteome, by assuming that cells have a fixed mass, *M*, at exponential growth. If the mass is dominated by the cell's proteins, then *M* is proportional to the size of the proteome in numbers of amino acids. At exponential growth, when the intracellular variables are at steady state, we can show (see SI Appendix, §S1.2.5) that if

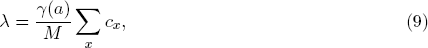

then

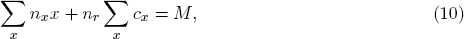

where *M* is approximately 10^8^ amino acids for *E. coli* [19] and assumed fixed (although *M* could also be made a function of *n_s_*, the quality of the available nutrients). Eq. 9 implements the trade-off through its enforcement of Eq. 10 (recalling that each *c_x_* contains a ribosome).

We assume that Eq. 9 holds generally and not just at exponential growth. The instantaneous growth rate is therefore the inverse of the time taken by the current number of translating ribosomes to synthesize all the proteins required for a new exponentially growing cell [21]. Although the mass of exponentially growing cells can vary with growth rate, we ignore such variations, which are typically small [19].

### The trade-offs capture fundamental properties of microbial growth

A model of exponentially growing microbes should recover general empirical properties of cell growth. The hyperbolic dependence of growth rate on levels of extracellular nutrients [15] is known as Monod's law and is a fundamental of microbiology. Two further relationships relate growth rate to the fraction of cellular mass comprising ribosomes: a linear, positive dependence as extracellular nutrients change (ribosomal mass fraction increases with growth rate) [16] and a linear, negative dependence as translation is inhibited by the addition of translation-poisoning drugs (ribosomal mass fraction decreases with growth rate) [2]. Although these growth relations have been observed in bacteria [22], there is some evidence that they are also valid in eukaryotes [23].

**Parameterizing the model:** We parameterize the model with parameters for *E. coli* from the literature (see SI Appendix, §S3) and then fit the remaining parameters to data from *E. coli* that demonstrate the two different types of linear dependence of ribosomal mass fraction on growth rate [2]. We fit parameters related to gene expression: the maximal transcription rates, *w_x_*; the transcriptional thresholds, *θ_x_* (Eq. 8); the auto-repression threshold for house-keeping genes, *K_q_*; and the translation threshold, *K_⋎_* (Eq. 3). In the experiments (Fig. 1 B),chloramphenicol was used to inhibit translation, and we model its action by having the drug sequester complexes of mRNA and ribosomes (see SI Appendix, §S3.1). We also therefore fit the rate constant for chloramphenicol binding, *k_cm_*.

The model fits the data of Scott *et al.* [2] (see SI Appendix, §S3.4, for a discussion ofthe quality of the fit) and reproduces the microbial growth laws (Fig. 1 B). No fine tuning of parameters is necessary: the model is robust in the sense that a range of parameters fits the data (Fig. 1 C& D and SI Appendix, §S3.3). We find that the transcriptional threshold for ribosomes, for in Eq. 8, is typically about two orders of magnitude larger than the transcriptional threshold, *θ_nr_*, used for all other genes with significant correlation (ρ = 0.85, *p*-value < 10^-20^; Fig. 1C & D). This difference in transcription thresholds implies that ribosomal and non-ribosomal transcription respond differently to cellular energy levels [4], and, as we shall see, this difference is key to allow the empirical growth relations to be derived from the model.

We emphasize that, although we parameterize our model with data from *E. coli*, the trade-offs considered are common to all growing cells, and so we expect the qualitative behaviour to be generally true. To apply specifically to another organism, the model should be re-fit to similar data.

**Deriving the growth relations:** The robustness of the model fit to the data suggests that the growth relations are an inherent property of the trade-offs comprised by the model. Indeed, under mild assumptions we can mathematically derive the relations from the model (see SI Appendix, §S2).

One relation is that growth rate is proportional to the ribosomal mass fraction, which follows from the definition of growth rate via ribosomal activity (Eq. 9) [2]. With *ϕ_R_* and *ϕ_r_* denoting the mass fractions of total and free ribosomes and τ_γ_ denoting the time for ribosomal synthesis, Eq. 9 can be rearranged to give (see SI Appendix,§S2.1)

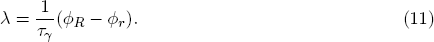

The synthesis time *τ_γ_ =n_r_ /γ* is the time taken to translate a ribosome and is a measure of ribosome efficiency: it relates the costs of ribosome production (the amount of energy required per ribosome) to the translational elongation rate. A smaller τ_γ_ implies higher ribosomal efficiency. Eq. 11 restates that the growth rate is proportional to the rate of translation and gives a linear dependence of the growth rate on the ribosomal mass fraction if τ_γ_ is approximately constant (for example, if the elongation rate γ(a) is near saturation at intracellular levels of a). Mechanistically, with more extracellular nutrient, more energy is available, which leads to more transcription. Transcription of ribosomes, however, is increased more than transcription of other proteins (θ_*r*_ ≫ θ_nr_ in Eq. 8), and so ϕ_*R*_ increases (Fig.2A).

Another empirical relation is a negative, linear dependence of the ribosomal mass fraction with growth rate when nutrient conditions are fixed and translation is inhibited by the addition of drugs (Fig. 1 B) [2]. We can derive

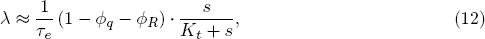

with ϕ_*q*_ being the mass fraction of non-ribosomal house-keeping proteins, *K_t_* being the Michaelis constant of the nutrient transporter, and τ_*e*_ being the enzyme synthesis time: the time taken to import sufficient nutrient to synthesize both a metabolic and a transporter enzyme. The enzyme synthesis time is therefore a measure of metabolic efficiency and is inversely proportional to the energy yield, τe ∼ *1/n_s_* (see SI Appendix, §S2.2). Eq. 12 therefore explains the different slopes obtained for different types of nutrients in Fig. 1B. Under the experimental conditions applied [2], we note that levels of extracellular nutrients, s, are constant, and so Eq. 12 is indeed linear. Intuitively, poisoning translation increases intracellular energy levels because fewer ribosomes can translate and leads to a proportionally larger increase in transcription of ribosomal mRNAs (θ_*r*_ ≫ θ_nr_) and so to a larger ϕ_*R*_. The negative dependence on ϕ_*R*_ arises because in this regime the growth rate is proportional to ϕ_*t*_, the mass fraction of the nutrient transporter, and so to the negative of ϕ_*R*_ because the total amount of proteins is conserved (see SI Appendix, §S2.2.3).

Finally, we can derive Monod's law to show a hyperbolic dependence of growth rate on the external nutrient s (see SI Appendix, §S2.3):

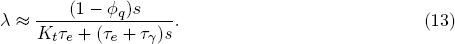

The maximal growth rate, 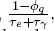, is determined by the mass fraction of non-ribosomal house-keeping proteins, φ_*q*_, and by the defficiency of ribosomes and metabolism. The half-maximal level of extracellular nutrients, 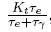, is proportional to the Michaelis constant of the nutrient transporter.

Our model recovers the growth relations because of both the trade-offs and the differences in transcriptional responses required by the data (θ_*r*_ ≫ θ_*nr*_). Several mechanisms can lead to differential transcriptional responses. For example, this difference could arise if RNA polymerases, whose levels increase with growth rate [19], have lower affinities to ribosomal genes either because of promoter structures or because the cell employs different polymerases for their transcription. Alternatively, in bacteria, ribosomal genes are enriched near the replication origin [24]. Consequently, the copy number of ribosomal genes will disproportionally increase through the parallel rounds of DNA replication used by bacteria during rapid growth [25] (when levels of energy are presumably higher), which can lead to increased levels of ribosomal transcription.

### Including the growth of the cell population

We can extend our model to include the growth of a population of cells (see SI Appendix, §S6.1). For a homogeneous population with a death rate of individual cells of *d_N_* ≥ 0, the number of cells, *N*, satisfies

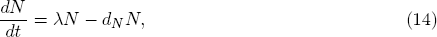

where the growth rate λ obeys Eq. 9. When all intracellular concentrations are at steadystate, the culture reaches exponential growth (SI Appendix, Fig. S5). The total amount of intracellular molecules in the population (across all cells) then grows exponentially.

By explicitly modelling the dynamics of extracellular nutrients, we can describe both batch and continuous cultures. For continuous culture, such as a chemostat, s has an influx rate *k_in_* and is diluted with a rate *d_N_* equal to the dilution rate of the cells. If each cell consumes nutrient with the same rate, *v_imp_*, we can describe the dynamics of the external nutrient by:

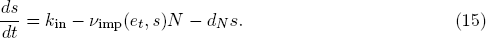

The steady-state number of cells is determined by the influx rate of nutrient and its energetic value *n_s_* and by the dilution rate and is approximately 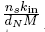 (see SI Appendix, §S6.1). For a batch culture, we set *d_N_ =k_in_ =0*, and consequently extracellular nutrient can only decrease from its initial amount. Eq. 15 then generates a typical growth curve with a lag phase if the number of nutrient transporters is, for example, initially low (SI Appendix, Fig. S5).

## Applications

### The trade-offs may explain gene dosage compensation for paralogs

With its parameterization from the data of Scott *et al.*, the model imposes global negative feedbacks on levels of enzymes and of ribosomes. Consider first the negative feedback on enzymes. If levels of enzymes fall, the cell imports and metabolizes less nutrient and energy levels decrease. Lower energy causes proportionally more enzyme mRNAs to be expressed (Fig. 2a) and consequently enzyme mRNA will be more successful in binding ribosomes. This success leads to increasing translation and so increasing levels of enzymes. Conversely, if levels of enzymes rise, energy levels rise and enzyme-mRNA will be less successful in binding ribosomes leading to decreasing levels of enzymes. The negative feedback on ribosomes works similarly. If levels of ribosomes fall, translation decreases and energy levels consequently rise causing proportionally more ribosomal transcription (Fig. 2a). An increase in ribosomes is in the same way counteracted by decrease in ribosomal transcription through changes in energy levels. The feedbacks act to balance energy in ux and consumption and so to stabilize energy levels.

Many genes have paralogs and the effects of deleting a gene can be reduced by increased expression of a paralogous gene, a phenomenon known as gene dosage compensation [17, 26]. Multiple global mechanisms can control gene expression [567]. For example, Keren *et al.* showed that the expression of most genes in both*E.coli* and *S.cerevisiae* is relative and stable at different growth rates [5]. We considered if dosage compensation could arise from the global coupling of gene expression and the negative feedback generated by the trade-offs comprising the model. For example, DeLuna *et al.* examined dosage compensation in over 200 genes in budding yeast and found that increased expression of a paralog upon deletion of its duplicated occurs only for genes required for growth [27].

**Figure 2.**
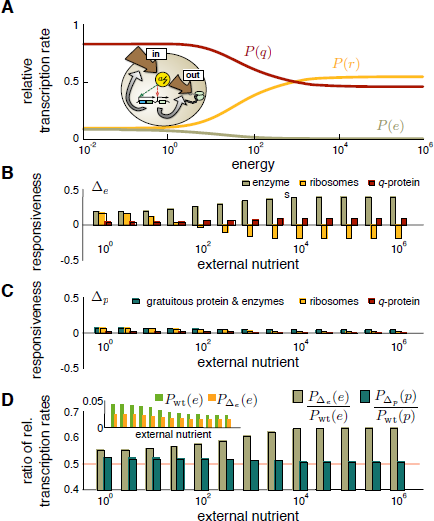
– The model can explain dosage compensation of pairs of paralogous genes. A. The relative abundance of mRNA changes with the level of intracellular energy because of different transcriptional responses of ribosomal and non-ribosomal genes (θ_*r*_ ≫ θ_*nr*_ in Eq. 8). We plot the relative transcription rate,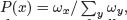, which determines the ability of *m_x_* to compete for ribosomes. Inset: schematic to illustrate the negative feedback via energy on levels of enzymes and ribosomes. **B.** Responsiveness is high upon deleting one of a pair of genes for an enzyme. Responsiveness in other genes is the log_2_ of the ratio of protein levels inthe deletion strain to those in the wild-type. For medium to high nutrient levels, ribosome-responsiveness is negative and so up regulating enzymes is at the cost of ribosomes. **C.** Responsiveness is low upon deleting one of a pair of genes for a gratuitous protein. **D.** Comparing the ratio of the relative transcription rates between the deletion and wild-type strains explains the corresponding behaviour of the responsiveness as a function of levels of external nutrient. A ratio above 0.5 (red line) implies dosage compensation. Inset: fractions of free enzyme-mRNA in the ▵e and the wild-type strains as a function of external nutrient.

To determine if this need-based regulation arises in the model, we first consider the deletion of an enzyme needed for growth and then the deletion of a gratuitous protein ― one that does not contribute to growth, but whose expression still uses global resources (see SI Appendix, §S4). Assuming that the paralogous gene copies are identical, we simulate a deletion strain, Δ_*x*_, by halving the maximal rate of transcription for a particular gene (*w_x_* in Eq. 8). For a system not constrained by cellular trade-offs and so with independent expression from each gene, levels of protein *x* in the deletion strain would be half the levels of protein *x* in the ‘wild-type’ strain where *wx* is unchanged. Dosage compensation occurs if these two quantities are not equal and can be quantified using the ‘responsiveness’ [27]: the log_2_ of the ratio of the levels of protein in the deletion strain to half the levels of protein in the wild-type strain. A system with independent gene expression would have a responsiveness of zero.

The model indeed predicts substantial dosage compensation for deletion of a gene for an enzyme, and the responsiveness increases with the level of available nutrients (Fig. 2B). Deleting a copy of the gene for an enzyme reduces the energy influx and so the steady-state levels of energy relative to the wild-type strain. The deletion strain correspondingly has proportionally higher levels of enzymes (Fig. 2A). The magnitude of the responsiveness as a function of external nutrients reflects an increase in the relative abundance of enzyme-mRNA compared to the wild-type strain (Fig. 2D). With high levels of nutrients, the transcription of enzyme-genes is saturated by the high levels of energy, but transcription of ribosomal genes still varies approximately linearly with energy (because θ_*r*_ ≫ θ_*e*_ in Eq. 8). Deleting an enzyme gene, which approximately halves the energy levels, reduces the rate of transcription of the enzyme genes, although not substantially (energy levels still exceed the transcriptional threshold θ_*e*_). The rate of transcription of ribosomal genes, however, halves. Reduced ribosome transcription relieves the competitive pressure for enzyme-mRNAs to bind ribosomes for translation, and so the frequency at which an enzyme-mRNA, rather than a ribosomal mRNA, succeeds in binding a ribosome is high. For low levels of nutrient, the rate of transcription of both ribosomal and enzyme genes varies approximately linearly with energy, and both are affected similarly by a reduction in energy levels. Consequently, the ratio between the relative transcription of enzyme-mRNA in the deletion and wild-type strains is low (and close to its theoretical minimum of 0.5).

Similarly, in agreement with DeLuna the model predicts little dosage compensation if we delete a copy of a gene for a gratuitous protein (Fig. 2C). Deleting a gratuitous gene affects energy levels substantially less than deleting a gene for an enzyme, and so the responsiveness is in general lower. In contrast to enzyme-deletion, deleting a gene for a gratuitous protein increases steady-state energy levels (although only by a few percent), and the responsiveness now decreases in high-nutrient environments, again following the trend in Fig. 2D. Unlike for enzyme-deletion, this latter behaviour does not re ect differences in energy levels because these differences are negligible. As levels of nutrients, and so levels of energy, increase, transcription becomes dominated by transcription of ribosomes. Hence the difference between whether the mRNA for the gratuitous protein is transcribed from one or two copies of the gene becomes negligible. The ratio of relative transcription of the mRNA of the gratuitous protein between the deletion and the wild-type strain tends to its minimum value of 0.5 (Fig. 2D).

In summary, the trade-offs that generate the growth laws also generate global negative feedbacks on proteins affecting growth. Whether this global regulation is the mechanism behind the observations of DeLuna however, requires further research: specific regulation, such as end-product inhibition of enzymatic pathways, is a possible alternative [27].

### Exploiting the trade-offs for host-aware design of synthetic circuits

A key goal in synthetic biology is to construct complex biochemical circuits with predictable functions [9, 28]. Synthetic circuits, however, compete for resources with their hosts in ways that are largely not understood. Host-circuit interactions can alter the designed function of a circuit [29], reduce the fitness of the host [8], and ultimately impose a negative selection pressure on cells with functioning synthetic circuits [30, 31]. Examples of competition effects include titration of native transcription factors [10]and cross-talk effects due to overloading of the degradation [32]or translation machinery [33].

**Figure 3.**
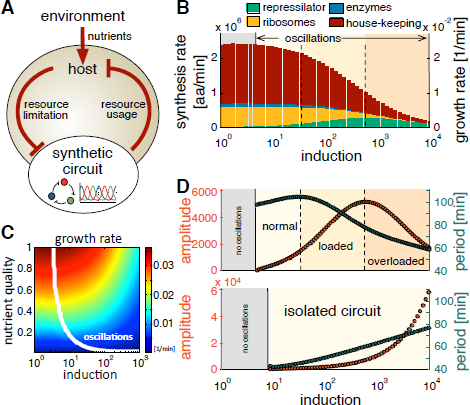
– The model predicts interactions between a synthetic circuit and its host cell. A. Aschematic of the interplay between the environment, host cell, and the repressilator as an example of a synthetic circuit. **B**.Growth rate and resources for host proteins decrease with increasing induction of the synthetic circuit. The strength of induction corresponds to the maximal rate of transcription of the repressilator genes. Stacked plots are the translation rates of different classes of proteins. Gray shading indicates induction levels where the levels of the repressilator proteins do not oscillate. Growth rate (linearly related to the total rate of translation - Eq. 9) equals the total height of the bars and is shown on the right hand axis. **C**.The range of induction needed for oscillations expands with a higher quality of nutrients and faster growth. The bifurcation curve between steady-state and oscillations is shown in white for different levels of induction and nutrient qualities. **D**. The repressilator behaves differently when simulated in isolation (lower panel) and within the cell model (upper panel): the host-aware model predicts a non-monotonic response that can be linked to loading of the host **(B)**

Our model can be used as a tool to quantify host-circuit interactions for the ‘host-aware’ design of synthetic gene circuits (Fig. 3A). The interplay between circuit, host, and environment can be directly incorporated into the design to minimize the impact of cellular trade-offs and resource competition on the circuit function. We can embed synthetic circuits in the model by defining new species linked to exogenous genes that compete for the shared pool of ribosomes and energy (see SI Appendix, §5). Although mathematical modelling is an integral part of synthetic biology's design cycle, most models do not include explicit interactions with the host[34]. These models cannot predict the impact of host-circuit interactions, resulting in an inefficient design process and lengthy trial-and-error iterations to appropriately tune a circuit's expression levels[35].

To illustrate the ability of the model to predict host-circuit interactions, we introduced a repressilator into the cellular chassis described by the model. The repressilator is a synthetic oscillator composed of three mutually repressive genes[36].The three repressilator proteins impose a burden on the cell, as they do not contribute to either growth or survival. To quantify the effects of host-circuit interactions, we focus on the impact of changing the levels of induction of the circuit (a commonly tuned quantity in synthetic circuits[37])and investigate growth and protein allocation in the host and the effect of changes in the host on the circuit's function.

The model predicts a sigmoidal decrease in growth for stronger induction of the repressilator genes (Fig. 3B). At low induction, expression of the synthetic genes is mostly at the expense of house-keeping proteins, including ribosomes. The host can compensate for this load and the consequent reduction of energy levels through transcriptional regulation and repartitioning of the proteome (following Fig. 2A). When the induction is sufficiently strong, however, competition for free ribosomes by the circuit mRNAs inhibits the synthesis of the host enzymes needed for nutrient transport and metabolism. This trade-off reduces expression of all proteins and consequently leads to a drop in growth.

We find that the onset of oscillations occurs at lower levels of induction as the growth rate increases (Fig. 3C). Since the oscillatory dynamics are driven by the negative feedback among the repressilator genes[36],the behaviour in Fig. 3C is likely to reflect a stronger negative feedback at faster growth rates because of higher numbers of repressor proteins. Fig. 3C provides a prediction of the model that can be directly tested by experiment. Further, the predicted behaviour suggests that environmental manipulations can be used to add flexibility to the design of synthetic circuits.

Host-circuit interactions can limit the ability to tune the behaviour of synthetic circuits. By comparing the function of the repressilator between the host-aware model and the traditional model isolated from the host (Fig. 3D), we observe significant differences in their oscillatory dynamics. The model of the isolated circuit predicts oscillations with amplitude and period that increase with the level of induction. The host-aware circuit, in contrast, predicts a non-monotonic behaviour because of over-loading of the host. For weak induction, and consequently little host-loading, the amplitude and period are qualitatively similar to those predicted by the isolated circuit, coinciding with a minor drop in growth (Fig. 3B). For intermediate induction, the period decreases with further induction and there is a major drop in growth. Once over-loaded, the amplitude too decreases reflecting an overall fall in protein production because of the limited synthesis of ribosomes (Fig. 3B). Further analysis suggests that such loading effects can be alleviated in environments richer in nutrients (SI Appendix, Fig. S4).

### Trade-offs can explain the evolution of gene regulation

Why one form of gene regulation has been selected over another is a fundamental question in both systems and evolutionary biology [38,39,40]. With our model's ability to link intracellular mechanisms to the growth of a cell population, we can investigate evolutionarily stable strategies by competing rival populations in silico. An evolutionarily stable strategy allows a population to resist invasion by any mutant population that uses an alternative strategy [41]. We consider the potential invasion of a resident population by mutant populations one at a time with deterministic simulations[42] (see SI Appendix, §6). The corresponding evolutionary assumptions, of weak rates of mutation and of large populations, are approximate and will not hold in general[43]

We let the maximum transcription rate of the enzymes be the evolvable trait (Fig. 4A and *w_e_* in Eq. 8) and model competitions between a resident strain with a particular *w_e_* and a mutant strain with a different value of *w_e_*. The resident population is allowed to reach steady-state in a chemostat (cf. Eq. 15) before a smaller mutant population appears. The two populations compete for the available nutrient and three outcomes are possible once the system reaches a new steady-state: (i) the mutant goes extinct and the resident resists invasion; (ii) the resident goes extinct and the mutant successfully invades; or (iii) neither the resident nor the mutant go extinct but both co-exist. By discretizing the range of values of we, we simulate all possible resident-mutant competitions and graphically show the results using invasion plots [42] (see Fig. 4B for an example)

First we consider growth in an environment with a constant influx of a single nutrient and find that the evolutionarily stable strategy is to have as high an expression of the enzymes as possible. We observe that a resident population with a maximal we is evolutionarily stable (Fig. 4C). The evolutionarily stable population has maximum expression of the transporter enzymes, reminiscent of the amplification of genes for transporters for nutrients limiting growth observed during adaptation in yeast [44]. Levels of enzymes are not tuned to match the availability of nutrients, but are always as high as possible to allow the population to outcompete any mutants. Growth of the resident population causes extracellular nutrients to fall until, at steady-state, each cell imports just enough energy to replicate over the time-scale determined by the dilution rate of a chemostat. A mutant with fewer transporters will be unable to import sufficient nutrient to match its growth rate to the chemostat's rate of dilution and will be lost.

This strategy although competitive is inefficient and generates a resident population with the smallest steady-state number of cells compared to resident populations with other values of the trait. Indeed, we see a rate-yield trade-off [45,46] (Fig. 4D), where a higher rate(proportional to the numbers of transporter enzymes) necessitates a lower yield (the numbers of cells in the population). This trade-off in rate versus yield at the level of the population is a consequence of the fundamental trade-offs in energy, free ribosomes, and proteins that act at the molecular level.

Regulated rather than constitutive expression appears almost universal. We postulated that a more nuanced strategy may arise when cells grow in environments with two nutrients because expressing genes to import and metabolize one nutrient will necessarily reduce expression of genes to import and metabolize the other. We therefore added to the model a second nutrient and a second set of constitutively expressed enzymes to import and metabolize that nutrient (Fig. 4E).

With a constant influx of two extracellular nutrients, an intermediate value of the maximum transcription rate can be evolutionarily stable, allowing the cell to balance the trade-off between exploiting one nutrient over another. Denoting the two nutrients by *s_a_* and *s_b_*, we let the maximum transcription rate for the *s_a_* enzymes, *w_a_*, be the evolvable trait and fix the maximum transcription rate for the *s_b_* enzymes. Invasion plots for different influxes of *s_a_* but a constant influx of *s_b_* are shown in Fig. 4F. When the influx of *s_a_* is lower than that of *s_b_*, the evolutionarily stable strategy is to minimize levels of the *s_a_* enzymes (*w_a_* is a minimum). The energetic cost of synthesizing the sa pathway is not compensated by the energy gained through metabolizing *s_a_*, and expression of the pathway is minimized (Fig. 4F left). Correspondingly, maximal cellular resources are freed for expression of the *s_b_* enzymes, suggesting that the competition for *s_b_* determines survival. In contrast, for a high relative influx of *s_a_*, we find that the evolutionarily stable strategy is to maximize levels of the *s_a_* enzymes (*w_a_* takes its maximum value: Fig. 4F right). Winning the competition for importing *s_a_* dominates, and the evolutionarily stable strategy maximizes expression of the *s_a_* transporters. For an intermediate influx of *s_a_* (Fig. 4F middle), competition for both nutrients determines if a mutant invades. An intermediate value of *w_a_* is evolutionarily stable, and this value increases with the influx of *s_a_* because of the greater importance of expressing sufficient *s_a_* transporters.

**Figure 4.**
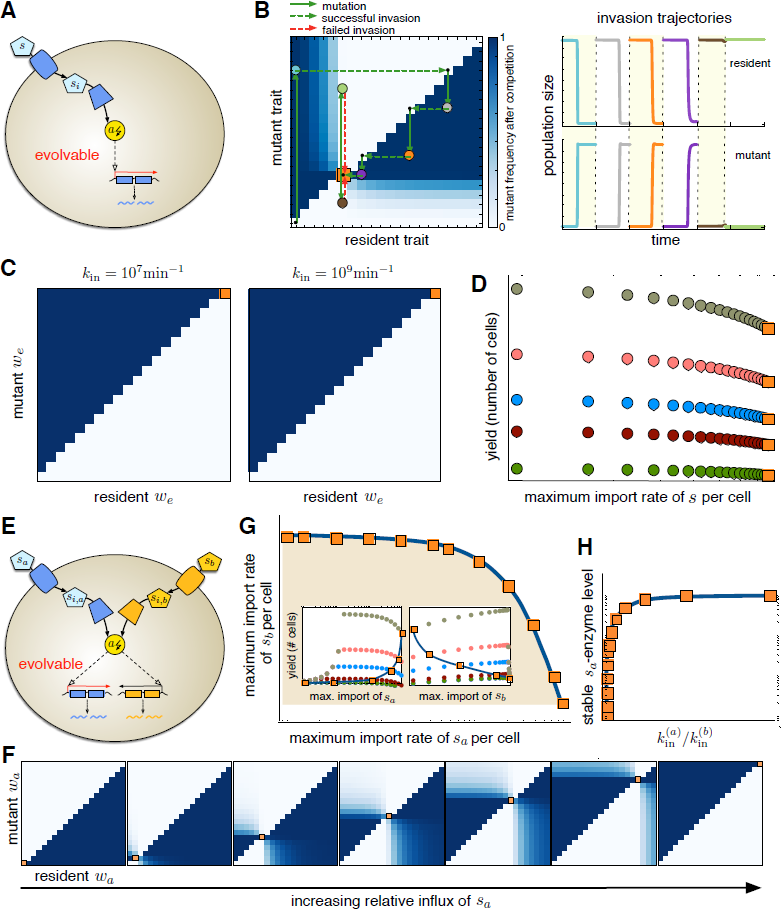
– Gene regulation is evolutionarily stable in changing environments because of a trade-off between metabolizing one type of nutrient over another. A. The maximum rate of transcription of the genes for the enzymes is an evolvable trait. **B**. An invasion plot (left) for an evolvable trait shows the results of all possible competitions between resident and mutant strains. The trait is assumed to take 20 discrete values and each square shows the result of one simulated competition. Colours indicate the steady-state mutant frequency 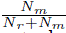 with *N_r_* and *N_m_* being the number of residents and mutants. White indicates that a mutant goes extinct; dark blue indicates that a mutant invades; light blue indicates co-existence. The result of a series of mutations is shown as a ‘cobweb’ plot, returning to the diagonal after each competition and repeating the process for the next mutation. Here the resident initially has a minimum value of the trait and is invaded by a mutant (blue dot). This mutant is itself invaded (gray dot), and the process repeats two more times (orange and purple dots). The evolutionarily stable value of the trait (orange square with white squares above and below) resists invasion of all possible mutants (two invasion attempts occur here). The simulations for the competitions for each mutation as a function of time are also shown (right). **C**. With a constant inux of a single nutrient, maximum expression of the enzymes is evolutionarily stable (*w_e_* is a maximum). **D**. The resident that has an evolutionarily stable *w_e_* (orange square) has a maximum rate of import of nutrients per cell (*v_t_ e_t_*, Eq. 7 in SI Appendix) but a minimum yield. All possible residents for five different influx rates of nutrient are shown (x-axis in log scale). **E**. With two nutrients, cells have two sets of enzymes, each specialized to import and metabolize one of the nutrients. Only the maximum rate of transcription of the enzymes for *s_a_* (*w_a_*) is assumed evolvable. **F**. Invasion diagrams for *w_a_* from models with two metabolic pathways show that an intermediate value can be evolutionarily stable. The influx of *s_a_* increases from left to right (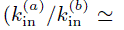 0.04, 0.7, 13, 55, 113, and 234 with 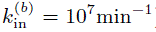). **G**. The steady-state enzymes levels are confined to a Pareto-like front (x-axis in log scale). The insets (similar to D) show that simple trade-offs in rate vs. yield do not exist: the evolutionarily stable values of *w_a_* can have low or high yields. H. The evolutionarily stable levels of enzymes for the *s_a_*-pathway as a function of relative influx rate suggest an evolutionarily stable strategy of regulation for changing environments (*y*-axis in log scale)

The steady-state dynamics is confined to a Pareto-like surface [47, 48] (Fig. 4G): maximal import of both *s_a_* and *s_b_* is impossible because trade-offs at the cellular level mean that increasing expression of one type of enzymes necessarily reduces expression of the other type. For different rates of influx of *s_a_*, the evolutionarily stable strategy moves on this surface reflecting the shifting importance of importing *s_a_* compared to *s_b_* as their relative abundances change, and we no longer see rate-yield trade-offs (Fig. 4G insets).

Our results point towards gene regulation being favoured in changing environments with multiple nutrients. For a single nutrient, the model suggests that constitutive expression should be selected because the evolutionarily stable strategy is to maximally express the enzymes regardless of environmental changes (modelled as changes in nutrient influx). With two nutrients, we see that constitutive expression is no longer evolutionarily stable, but instead that the expression of the *s_a_* enzymes should be regulated (and follow the relation in Fig. 4H).

## Discussion

By constructing a model based around three fundamental trade-offs that are faced by all living cells in their use of energy, ribosomes, and mass, we have shown that we can explain both empirically derived growth relations for bacteria and potentially dosage compensation by paralogs in budding yeast. Further, our model predicts the effects of similar trade-offs generated by synthetic circuits in host cells and can be extended to include the growth of cell populations.

We have adopted a coarse-grained approach to increase the generality of the model and to highlight basic mechanisms driving phenotypic change, but our model can be extended in multiple ways. For example, explicit mechanisms for the dependence of both transcription on energy and translation on levels of tRNAs, which are known to change with growth rate [49, 50], could be included. Such additions, however, lead towards whole-cell modelling [11], and our approach has been to try to include the minimal biochemistry necessary to answer the questions of interest. Our framework could be adapted to describe different organisms by, ideally, changing parameter values while a whole-cell model is inherently specific to a particular cell type.

Through its coupling of biochemistry to growth rate to populations, the modelling framework we propose has several immediate translational applications. First, many antibiotics target dividing cells. By including the action of these antibiotics in the model, we should be able to predict the effects of suppressive drug interactions [51], where one drug can ameliorate the consequences of another, and of any feedback between growth rate and gene expression generated by antibiotics affecting translation [52]. Second, we have illustrated how to predict trade-offs between the induction level of a synthetic circuit, its function, and the growth of the host. We can therefore benchmark different designs aimed at producing chemicals in biotechnology, where circuits must operate robustly in different growth conditions [53, 54]. Third, disregulated biogenesis of ribosomes has been suggested as a driver for cancer development [55], and our model may help select, for example, therapeutic targets in the translation machinery.

Genes are not expressed in isolation but through all stages of expression interact with the surrounding molecules that comprise living cells. These interactions create the potential for trade-offs and including such aspects of cell physiology has great promise for predicting phenotypic quantities from genotypic specifications, a long-term goal of both systems and evolutionary biology.

## Methods

**Simulations:** Details of all model assumptions and equations (Eqs. 1-10) along with the parameter values taken from the literature is given in the SI Appendix (§S1). SBML and Matlab versions of the model are also available. To simulate the model, we used ode15s from Matlab's stiff integration suite. For the synthetic gene circuit, we adapt the original repressilator model [36], adding equations for the three synthetic proteins, together with their free and ribosome-bound mRNA, to the model and modify the energy and ribosome usage and the growth rate accordingly (see SI Appendix, §S5.1). To study the dynamics of competing strains, we duplicate all model variables, except those for extracellular nutrients, to describe the resident and mutant populations and include consumption of nutrients by both populations (see SI Appendix, §S6).

**Parameter fitting:** To fit the undetermined parameter values to the data from Scott [2], we used a Bayesian approach with an adaptive Markov chain Monte Carlo sampling procedure [56]. We simulated the model for various (fixed) nutrient quality values (*n_s_*) at the given concentrations of chloramphenicol to predict growth rates and the fractions of ribosomal protein mass at steady-state and so calculate the likelihood of the parameters given the data. The final parameter values chosen correspond to the modes of the marginal posterior distributions. From the posterior distribution we further estimated the Fisher information matrix and parameter sensitivities [57], which indicated a robust fit to the data (SI Appendix, Fig. S2).

## Acknowledgments

We thank Ivan Clark, Ramon Grima, Nick Jones, Nacho Molina, Vahid Shahrezaei, Guillaume Terradot, Philipp Thomas, and particularly Ted Perkins and Matt Scott for helpful comments and suggestions. AYW is grateful for support from a DFG Research Fellowship; DAO acknowledges support from an Imperial College London Junior Research Fellowship; VD from the ERC; and PSS from the Scottish Universities Life Sciences Alliance.

